# Disruption of c-MYC binding and chromosomal looping involving genetic variants associated with ankylosing spondylitis upstream of *RUNX3* promoter

**DOI:** 10.1101/2021.09.07.459234

**Authors:** Carla J Cohen, Connor Davidson, Carlo Selmi, Paul Bowness, Julian C Knight, B Paul Wordsworth, Matteo Vecellio

## Abstract

**Background:** Ankylosing Spondylitis (AS) is a common form of inflammatory spinal arthritis with a complex aetiology and high heritability, involving more than 100 genetic associations. These include several AS-associated single nucleotide polymorphisms (SNPs) upstream of *RUNX3*, which encodes the multifunctional RUNT-related transcription factor (TF) 3. The lead associated SNP *rs6600247* (p= 2.6 x 10^-15^) lies ~13kb upstream of the *RUNX3* promoter adjacent to a c-MYC TF binding-site. The effect of *rs6600247* genotype on DNA binding and chromosome looping were investigated by electrophoretic mobility gel shift assays (EMSA), Western blotting-EMSA (WEMSA) and Chromosome Conformation Capture (3C).

**Results:** Interrogation of ENCODE published data showed open chromatin in the region overlapping *rs6600247* in primary human CD14+ monocytes in contrast to Jurkat T cell line or primary T-cells. The *rs6600247* AS-risk allele is predicted to specifically disrupt a c-MYC binding-site. Using a 50bp DNA probe spanning *rs6600247* there was consistently less binding to the AS-risk “C” allele of both purified c-MYC protein and nuclear extracts (NE) from monocyte-like U937 cells. WEMSA on U937 NE and purified c-MYC protein confirmed these differences (n=2; p<0.05). 3C experiments demonstrated negligible interaction between the region encompassing *rs6600247* and the RUNX3 promoter. A stronger interaction frequency was demonstrated between the *RUNX3* promoter and the previously characterised AS-associated SNP *rs4648889*.

**Conclusions:** The lead SNP *rs6600247*, located in an enhancer-like region upstream of the *RUNX3* promoter, modulates c-MYC binding. However, the region encompassing *rs6600247* has rather limited physical interaction with the promoter of *RUNX3*. In contrast a clear chromatin looping event between the region encompassing *rs4648889* and the *RUNX3* promoter was observed. These data provide further evidence for complexity in the regulatory elements upstream of the *RUNX3* promoter and the involvement of *RUNX3* transcriptional regulation in AS.

## BACKGROUND

Ankylosing Spondylitis (AS) is a form of inflammatory spondyloarthritis predominantly affecting the axial skeleton, which is characterised pathologically by enthesitis (1). Extra-skeletal manifestations are also common in AS; these include inflammation of the gut (ranging from low-grade sub-clinical inflammation of the terminal ileum to overt inflammatory bowel disease - IBD), skin (psoriasis), and uveal tract (2, 3). AS was one of the first complex diseases in which a specific genetic effect was identified when its strong association with the major histocompatibility complex (MHC) immune response gene HLA-B27 was described nearly fifty years ago (4) (5). However, it is clearly polygenic (6); even the MHC association is attributable to several alleles at more than one locus (7) and more than 100 non-MHC genetic associations have now been suggested by genome-wide association studies (8)(9)(10)(11). Shared genetic susceptibility factors undoubtedly contribute to the excess occurrence of psoriasis, IBD and uveitis not only in individuals with AS but also their relatives (12) (13). One of the strongest non-HLA associations with AS is with the *RUNX3* (Runt-related transcription factor (TF) 3) locus. RUNX3 is involved in T-cell function and plays a key role in the development of CD8+ T-cells.(14) It also influences many other cells, including helper T-cells, innate lymphoid, tissue resident, mucosa and gut cells (15, 16). We have recently demonstrated that AS-associated non-coding single nucleotide polymorphisms (SNPs) in an enhancer-like region upstream of *RUNX3* affect the binding of different factors: in particular the repressive nucleosome remodelling and deacetylase (NuRD) complex binds preferentially to the risk allele, while conversely interferon regulatory factor (IRF) 5 to the protective allele (17). However, the functional effects of these changes on gene transcription are still to be precisely determined. Our earlier observations were made in T-cells, but here we describe some of the functional effects of the lead AS-associated SNP in the vicinity of *RUNX3* (*rs6600247*, p = 2.6 x 10^-15^ (11) that are more obvious in CD14+ monocyte-like cells than CD8+ T-cells. First, we evaluate the chromatin landscape surrounding *rs6600247* using the ENCODE database (https://genome.ucsc.edu/ENCODE/). Second, we demonstrate differential allelic binding of *rs6600247* to the c-MYC TF. Finally, we investigate the chromosomal architecture and physical interactions between AS-associated sequences in the enhancer-like region upstream of *RUNX3* and its promoter, showing a probable role of chromosome looping in the regulation of *RUNX3*.

## METHODS

### Genotyping

DNA was extracted using the Qiagen AllPrep DNA/RNA Mini Kit (Qiagen Ltd, Manchester, UK) and genotyped for *rs6600247* using TaqMan SNP assay (custom order by Life Technologies, Paisley, UK), for the cells (obtained by the buffy coat) used in the functional studies.

### *In silico* investigation

We used the UCSC genome browser build hg19 and the Roadmap database [https://genome.ucsc.edu/ENCODE/] to investigate the epigenetic landscape of *rs6600247* upstream of the *RUNX3* promoter, which is strongly associated with AS (p=4.2×10^−15^) (11). Histone modifications and GeneHancer (a database of human regulatory elements and their inferred target genes) tracks were selected to evaluate regulatory elements and chromosome looping between promoters and enhancer regions (18).

### Cell lines, culture and primary human cell isolation

CD8+ T-cells and CD14+ monocytes were isolated from AS patients’ peripheral blood mononuclear cells (PBMCs) using a CD8+ T-cell or a CD14+ monocyte isolation kit (Miltenyi, Bisley, Surrey, UK), respectively. Jurkat, U937, CD8+ and CD14+ cells were resuspended at 1×10^6^/mL in pre-warmed Roswell Park Memorial Institute medium supplemented with 10% fetal bovine serum, penicillin/streptomycin and L-glutamine, and rested overnight. Cells were then harvested for experiments.

### Electrophoretic mobility gel shift assay (EMSA)

The impact of *rs6600247*, which lies in a c-MYC binding-site (Figure 2A), was assessed by EMSA. We designed DNA probes including either the protective T or the AS-risk variant C to evaluate the disruption of a C-MYC consensus motif. The DNA probes used in EMSAs (50-bp single-stranded biotinylated DNA probe incorporating *rs6600247*) were mixed and annealed at room temperature for 1□hour. Probes were then incubated for 20 min. with nuclear extracts (NE) obtained either from primary CD8+ T-cells or a monocyte cell line from histiocytic lymphoma (U937) stimulated with phorbol-12-myristate-13-acetate (PMA). The sequences of the synthetic single-stranded oligonucleotides are listed below:

C* s (sense): 5′-CTCCATGACGCAATTTGGGCTC<u>C</u>GTTATGAGTCAGCTCAAGTAA-3′; T* s: 5′-CTCCATGACGCAATTTGGGCTC<u>T</u>GTTATGAGTCAGCTCAAGTAA-3′;
C* as (antisense): 5′-TTACTTGAGCTGACTCATAAC<u>G</u>GAGCCCAAATTGCGTCATGGAG-3′; T* as: 5′-TTACTTGAGCTGACTCATAAC<u>A</u>GAGCCCAAATTGCGTCATGGAG-3′.

(Underlined base highlights the position of *rs6600247*).

### Western Blotting - Electrophoretic mobility gel shift assay (WEMSA)

DNA probes as for EMSA (above) were incubated with nuclear extract obtained from U937, CD8+ T-cells or purified c-Myc human recombinant protein (Abcam, ab169901 Cambridge, UK) as previously described (19) and separated on DNA retardation gels at 100 V on ice. The samples were transferred on to nitrocellulose membranes for Western blotting (WB), then blocked with 5% milk in Tris Buffer Saline + 0.1% Tween (TBST) for 1 hour at room temperature (RT) before incubating overnight at 4°C with the primary antibody for c-Myc (Santa Cruz Biotechnology sc-40, Dallas, Texas USA).Secondary goat anti-rabbit antibody (1:10000 dilution) was added (1hour RT) and the membranes washed before Horse Radish Peroxidase substrate (Thermo Fisher Scientific, Waltham, Massachusetts, US) added for imaging. Image J (NIH) was used for quantifying WEMSA bands (20).

### Chromosome Conformation Capture

Chromosome conformation capture (3C) was performed as previously described (21). Briefly, libraries were prepared as follows: 1.5×10^7^ of U937 or Jurkat cells were cross-linked with formaldehyde at 1% of the final volume. Glycine [0.125M] was used to quench cross-linking and cells were lysed in cold lysis buffer on ice using a Dounce homogenizer (Sigma Aldrich, Gillingham, UK). Cells were resuspended in specific restriction enzyme buffer (10 μL were kept as undigested control). The remaining samples were digested overnight at 37°C with 500 units of SacI (New England Biolabs, Hitchin, UK). Digestion was stopped by the addition of 10% sodium dodecyl sulfate incubated at 65°C for 30 min. T4 ligase (Ambion, Thermo Fisher Scientific, Waltham, Massachusetts, US) was used to perform ligation for 4 hours at 16°C. Proteinase K was added prior to reversal of cross-linking at 65°C overnight. Proteinase K was added to the undigested and digested controls saved earlier. DNA was purified using phenol-chloroform extraction, followed by ethanol precipitation. 3C template was resuspended in 500 ul H_2_O, while undigested and digested controls in 50ul. The quality of the chromatin samples was assessed on agarose gels. Bacterial Artificial Chromosome (BAC) preparations were performed similarly as genomic controls. 3C PCR primers were designed along the same strand and same orientation to accomplish specific amplification across 3C ligation junctions. Full list of primers is available in Supplementary table 1 and their genomic position relative to *RUNX3* is shown in Figure 3

We interrogated a genomic region upstream the *RUNX3* distal promoter, including few AS-associated SNPs in U937 (monocyte-like) and Jurkat (T-lymphocyte-like) cell lines. The bait was placed at the distal promoter (P2) with amplification primers at the AS-associated SNPs *rs6600247* and *rs4648889* along with three intergenic regions.

### Quantitative real-time polymerase chain reaction (qRT-PCR)

Total RNA from CD8+ and CD14+ cells was isolated with TRIzol (Invitrogen, Paisley, UK) and reverse transcribed with Superscript III (Invitrogen, Thermo Fisher Scientific, 168 Third Avenue, Waltham, Massachusetts, US) to synthesise cDNA as previously described. (22) The specific primers were: RUNX3 sense (s): 5′-ACTCAG CAC CAC AAG CCA CT-3′; RUNX3 antisense (as): 5′-GTC GGA GAA TGG GTT CAG TT-3′. Quantitative PCR was performed in triplicate and the 2−ΔCt method was used to calculate the expression of *RUNX3* relative to β-actin (ID Assay qHsaCED0036269, Bio-Rad Laboratories, Kidlington, UK).

### Historical controls and RUNX3 expression

RUNX3 transcription in AS cases and controls was evaluated from previously published data derived from RNA-seq in PBMCs from 72 AS cases and 62 healthy controls and stratified for *rs6600247*. (23)

## RESULTS

### Genomic landscape interrogation suggests a regulatory role for *rs6600247*

Figure 1A shows the genomic landscape at the *RUNX3* locus, with the lead AS-associated SNP *rs6600247* lying ~13kb upstream of the distal promoter while the regulatory SNP *rs4648889* is physically closer to the promoter. SNP *rs6600247* is situated within a region of open chromatin, defined by a peak for DNase I hypersensitvity (DHS - indicative of regions of open chromatin) (Figure 1B), and a peak of H3K4Me1 histone modification. This sequence also binds the transcription factor c-MYC (ENCODE Factorbook (http://www.factorbook.org/human/chipseq/tf/) (Figure 1B). Taken together, these data suggest an enhancer-type element surrounding *rs6600247*, so we sought to determine a regulatory role of this SNP.

**Figure 1.**
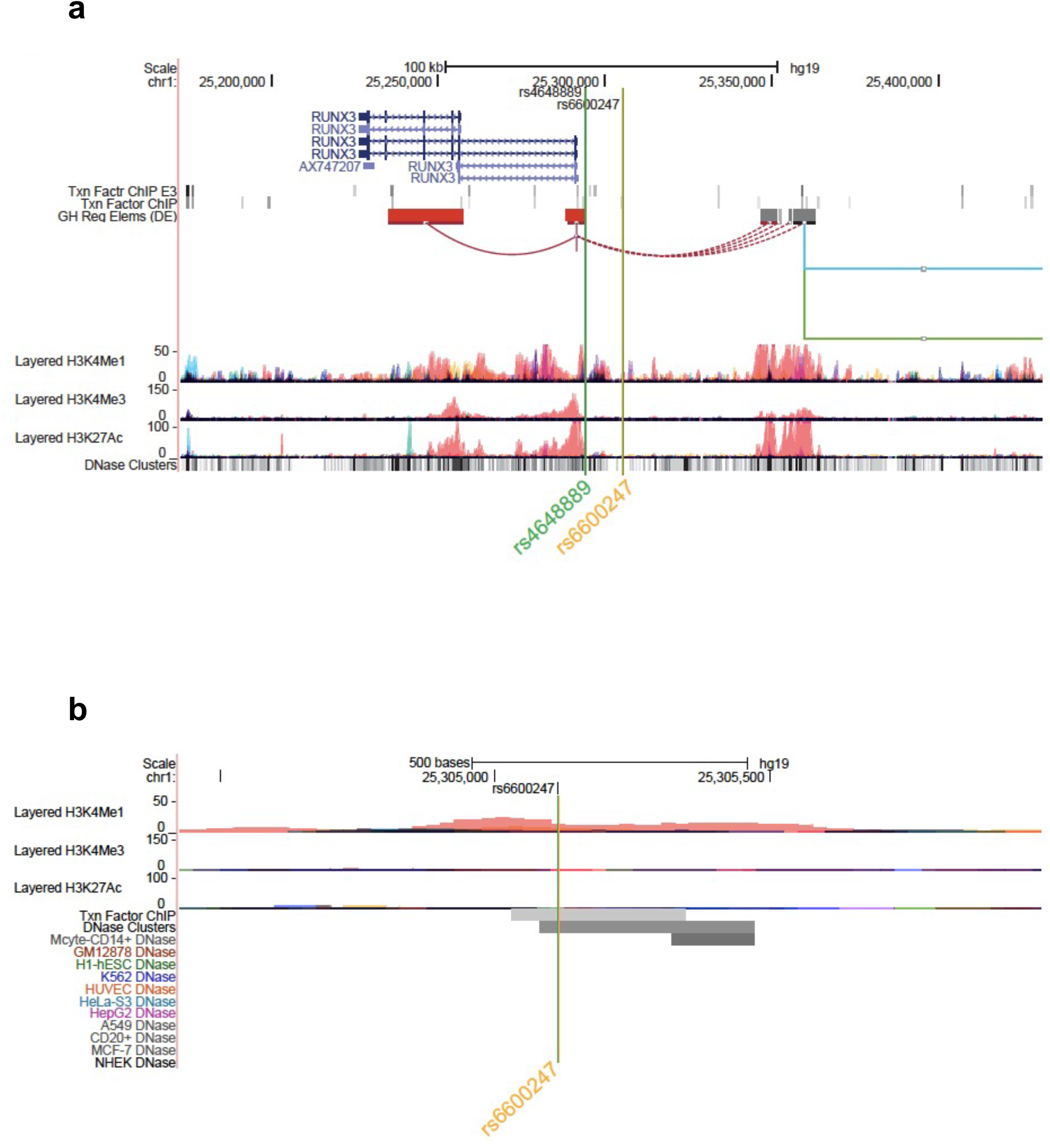
Genomic landscape interrogation suggests a regulatory role for *rs6600247*. UCSC Genome Browser analysis of the *RUNX3* locus at chr1:25,172,487-25,431,044. The green and orange vertical lines show the location of SNP *rs4648889* and *rs6600247*. *RUNX3* gene location (blue) is from the UCSC genes database. Two tracks show transcription factor binding by ChIP (Txn Facto ChIP E3 and Txn Factor ChIP). GeneHancer track shows human regulatory elements and their inferred target genes. ENCODE layered H3K27ac, H3K4Me1 and H3K4me3 tracks (from seven cell lines) show enrichment of these marks, indicating promoter and enhancer regions. DNase clusters show DNase-I hypersensitivity clusters (ENCODE); B) Zoomed Genome Browser view (chr1:25,304,328-25,305,991) upstream of the promoter of *RUNX3* showing a peak for H3K4Me1 enrichment overlapping *rs6600247* (vertical line; Layered tracks as in A). Additional ENCODE datasets displayed are transcription factor Chip-seq peaks and DNaseI HS peaks that directly overlap *rs6600247*,and DNase I HS for CD14+ monocytes in proximity to *rs6600247*.

The DHS peak overlapping *rs6600247* is seen specifically in CD14^+^ monocytes (Figure 1B). For this reason, we conducted our functional experiments in U937 cells, a pro-monocytic, human myeloid leukaemia cell line, exhibiting monocyte-like features.

### *rs6600247* AS-risk C allele alters c-MYC binding to DNA

We analysed the DNA sequence at *rs6600247* and found that the SNP lies within a c-MYC consensus binding motif (Figure 2A). We hypothesised that binding of c-MYC protein to a DNA sequence containing the risk allele C would be reduced. The results of EMSA assessing the relative c-MYC protein binding to the C or T alleles are shown in Figure 2. We first incubated probes with recombinant c-MYC purified protein, and observed a specific DNA/protein complex with both alleles but markedly less to the AS-risk allele C than the protective T allele. (Fig 2B; lane 3-4, n=3). We then incubated the same probes with NE from U937 (monocyte-like cells) and observed a major protein/DNA complex binding to the protective T allele, but none with the C allele (Fig 2C, lanes 3-4, n = 3). In both cases, successful competition with a 100-fold excess of unlabelled probe confirmed the specificity of the complex (Fig 2B, lane 5-6 and Fig 2C lane 5-6). We next used WEMSA to quantitate the relative binding of c-MYC to each allele of *rs6600247*. Markedly less c-MYC enrichment was seen with the C risk vs T allele using either c-MYC purified protein or U937 NE (Fig 2D and 2E, relative band intensities p=0.01 and p=0.05, respectively, two-sample *t* test).

**Figure 2.**
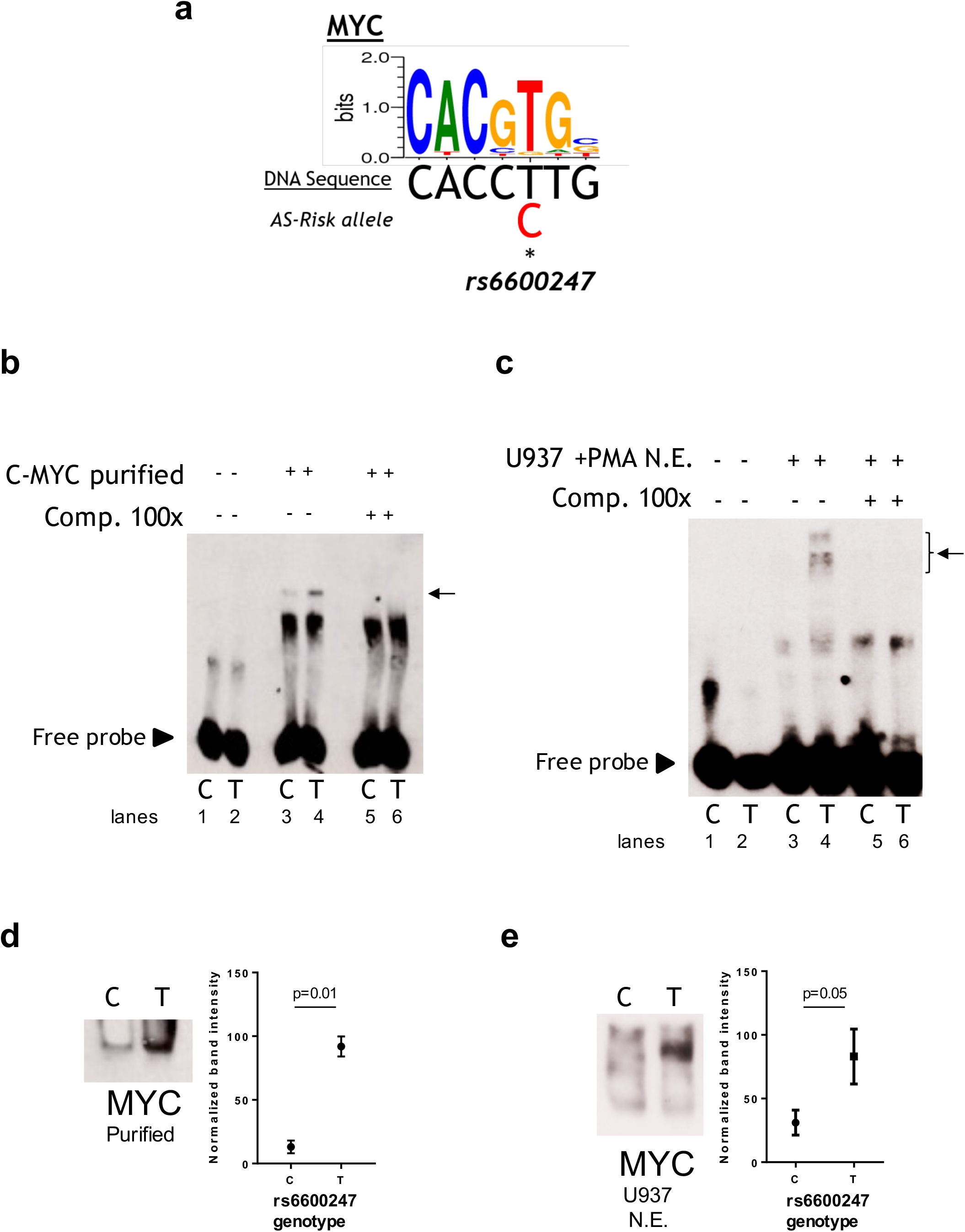
*rs6600247* risk allele affects C-Myc binding. A) C-Myc binding motif analyzed using the MEME program and the location of *rs6600247* risk allele; B) EMSA using c-MYC purified protein with or without specific competitor (Comp 100x), n=4, allele (C or T) of *rs6600247* included in the 50bp biotinylated double-stranded DNA probe is given below the image; horizontal arrow indicates specific protein-DNA complex formation; C) EMSA using nuclear extract (N.E.) from U937 cells stimulated with phorbol 12-myristate 13-acetate (PMA) with or without specific competitor (Comp 100x), n=4; *rs6600247* allele and complex formation indicated as in B; D) WEMSA using C-Myc purified protein and blotted with an antibody against C-Myc; E) WEMSA using U937 nuclear extract and blotted with an antibody against C-Myc. The blot is representative of n=2 experiments. Binding in D) and E) was quantified using ImageJ software and is representative of two different experiments, demonstrating that the risk allele for rs6600247 shows fewer binding properties for C-Myc.

### The *RUNX3* promoter interacts with the *rs4648889* region rather than *rs6600247*

We used 3C to test plausible chromosome looping interactions between the AS-associated SNP *rs6600247* and the *RUNX3* distal promoter. Figure 3A shows the RUNX3 genomic region interrogated, the location of the primers and Sac I restriction sites. Baits were designed to capture SacI fragments containing *rs6600247*, three intergenic fragments with H3K4me1 enrichment, and additionally with a previously-studied AS-associated SNP *rs4648889*. There was very low interaction frequency between *rs6600247* and the distal *RUNX3* promoter, either in U937 or Jurkat cells. A stronger interaction frequency was observed between the distal promoter and the region encompassing the AS-associated SNP *rs4648889* (Figure 3B) confirming its functional role.

**Figure 3.**
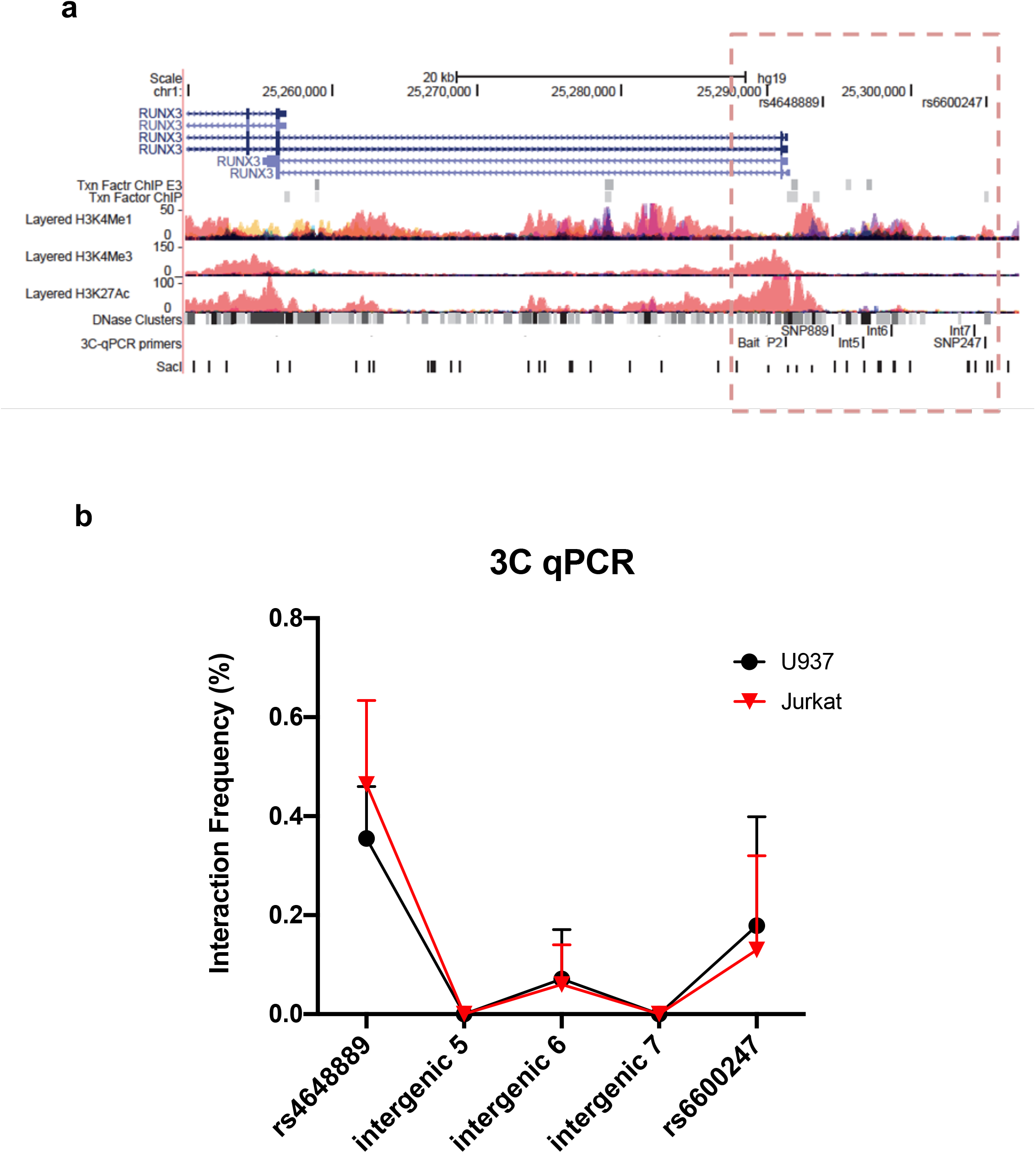
Chromosome looping investigation demonstrates interaction between the *RUNX3* promoter and region encompassing *rs4648889* rather than *rs6600247*. A) Location of the *RUNX3* genomic region chr1:25,249,544-25,307,338. Tracks shown as in Figure 1, with the addition of 3C-qPCR primers and SacI enzyme cutting sites. Dotted box highlights the region analysed to identify potential amplified interaction fragments tested with 3C. Bait fragment is located at *RUNX3* distal promoter (P2); AS-associated SNPs primers used in these experiments are named as follows: SNP889 (for *rs4648889*), Int5, Int6, Int7 (for intergenic regions 5, 6 and 7) and SNP247 (for *rs6600247*); B) Results of the 3C-qPCR analysis showing Increased relative interaction frequency between *RUNX3* P2 and the region encompassing *rs4648889*, with a modest interaction with the *rs6600247* region. Theses interactions were seen in both U937 (red) and Jurkat (black) cell lines.

### *rs6600247* genotype has no effect on *RUNX3* expression

Primary CD14+ monocytes and CD8+ T-cells from AS patients were used to evaluate *RUNX3* mRNA expression stratified on *rs6600247* genotype (n=5 each genotype) (Figure 4A and 4B). There was a non-significant trend for lower expression in CD14+monocytes with the AS-risk CC genotype compared to protective TT and heterozygous TC genotypes (TT vs CC: 4.6±1.8 vs 2.0±0.4; TT vs CT: 4.6±1.8 vs 1.8±0.2; CC vs CT: 2.0±0.4 vs 1.8±0.2, results are expressed as mean ± standard error mean). We also analysed historical RNA-seq data (23) obtained from AS patients’ PBMCs measuring *RUNX3* mRNA expression, stratified on *rs6600247*: there was no apparent influence from this SNP on *RUNX3* expression (Figure 4C).

**Figure 4.**
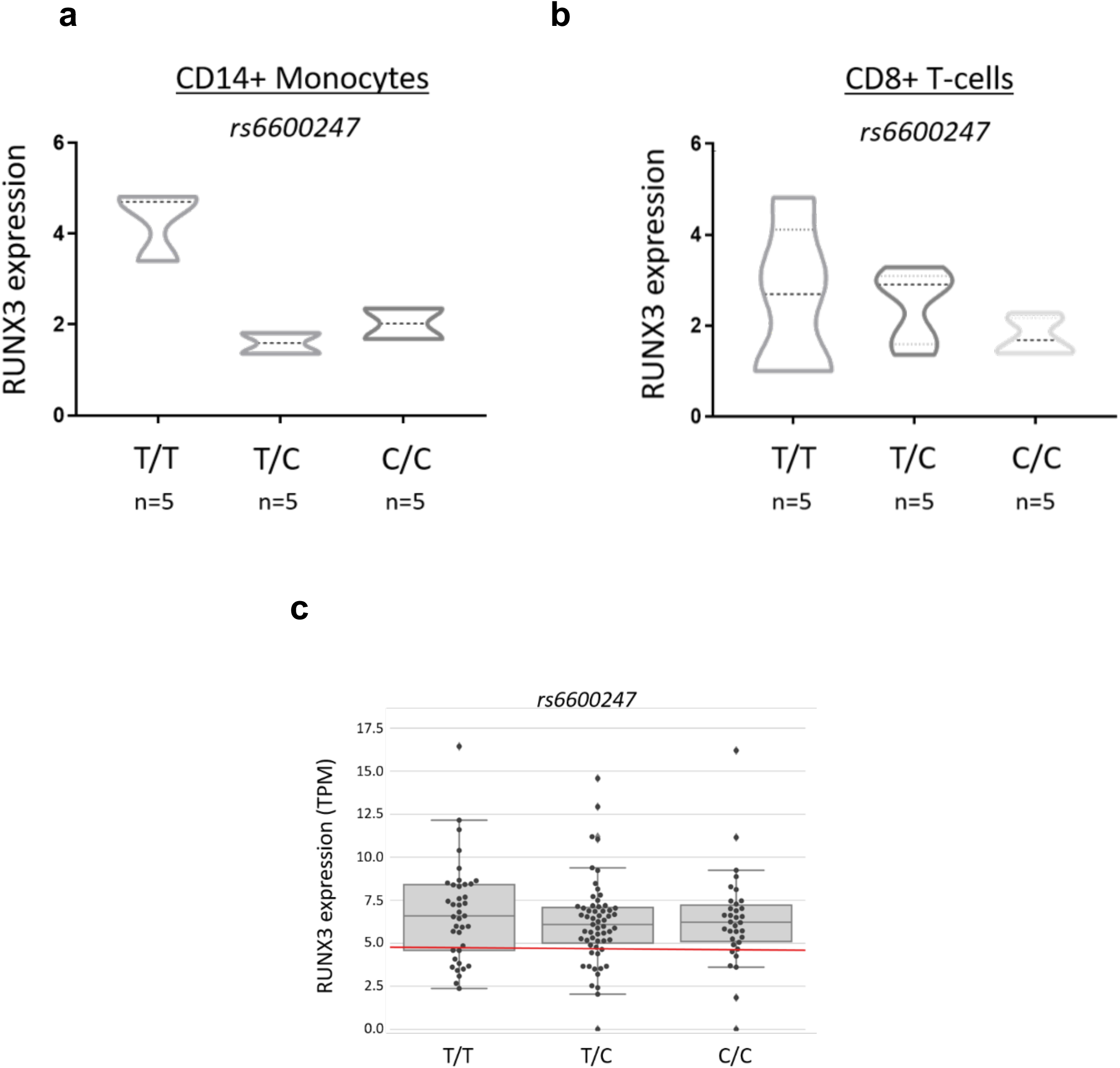
*rs6600247* genotype shows no regulatory effect on RUNX3 expression. *RUNX3* expression levels measured by qRT-PCR in freshly isolated A) CD14+ monocytes and B) CD8+ T-cells from 15 AS patients stratified according to *rs6600247* genotype. Statistical analysis performed with Welch’s two-sample *t* test; C) Expression of *RUNX3* in an historical RNA-seq dataset (23) obtained from PBMCs, stratified on *rs6600247*.

## DISCUSSION

In this study, we have demonstrated that the AS-associated SNP, *rs6600247*, affects the binding of c-MYC to this region of DNA, 13 kb upstream of the *RUNX3* promoter, which lies in a region of open chromatin in CD14+ monocytes. Although GWAS have identified hundreds of genetic variants associated with AS, (11) only a very small portion of these have been investigated to define causal variants. Cell type and stimulation conditions must be taken carefully in consideration in identifying causal SNPs, as both impact on chromatin interaction and gene regulation. (24)

Recent findings have shown that RUNX3 is highly expressed in monocytes where it has a role in transcriptional repression, metabolic regulation, and in tuning the function of CD14+ monocytes. (25, 26). Expression of RUNX3 has been found also in CD11c+ mature intestinal macrophages, suggesting a role for this TF in macrophage maturation.(27) Further, both RUNX3 and another TF, ID2 (Inhibitor of DNA binding 2), are required for the differentiation of epidermal Langerhans cells from monocytes. (28)

The interaction between RUNX3 and c-MYC has previously been investigated in T-cell lymphoma.(29) Double immunofluorescence revealed co-localization of both proteins in the tumour nuclei. In addition, several binding-sites for c-MYC were identified in the RUNX3 enhancer region. Additional evidence for this interaction stems from colorectal cancer studies where upregulation of RUNX3 by Bone Morphogenetic Protein (BMP) reduces c-MYC expression, thereby exerting c-MYC tumour-suppressor activity.(30) Recently, it has been demonstrated that two superenhancers located at 59 and 70 kb upstream of the RUNX3 transcription start site are required for both RUNX3 and MYC expression and function.(31) Other studies also indicate a key role of c-MYC in monocyte/macrophage activation, as it is involved in the regulation of different alternative activation genes.(32)

Our EMSA/WEMSA experiments were able to confirm c-MYC binding the *rs6600247* locus, with the AS-risk allele disrupting the binding motif and consequently reducing formation of the c-MYC-DNA complex. Altogether, these observations are consistent with the hypothesis that c-MYC can bind the *RUNX3* promoter and/or regulatory elements upstream of the promoter thereby playing a role in the functional regulation of *RUNX3*. The processes involved in transcriptional regulation are complex and this finding does not exclude the possibility of other TF being involved.

The genome is organized in a very dynamic way and TFs mediate chromosome loops to bring enhancers and promoters together. (33, 34) 3C and related techniques are the paradigmatic approach to demonstrate interactions between target genes and enhancers or enhancer-like regions. Here, we have demonstrated physical interaction between the distal promoter of *RUNX3* and *rs4648889* SNP, which we have previously functionally characterized by our group. (17, 35) Conversely, a very low interaction was observed with *rs6600247* suggesting no evident functional role for this SNP in chromosome looping in this particular cellular context.

It is now clear that the presence of an enhancer-promoter loop alone does not ensure activation of a target gene: this provides a platform where transcription factors can bind and regulate gene/s.(36, 37) Here, we have confirmed that the genomic regulatory element upstream of the RUNX3 promoter has potentially important cell-type-specific functional effects. We showed that the AS-risk allele of *rs6600247* affects c-MYC binding in monocytes, suggesting c-MYC/RUNX3 pathway should be further investigated for its potential role in the pathophysiology of AS. Nevertheless, the region encompassing *rs6600247* has no significant physical interaction with the distal *RUNX3* promoter, confirming *rs4648889* the cardinal genetic variant associated with AS of major interest within the RUNX3 genomic locus.

Further studies are required to better elucidate the presence of additional higher order chromatin interactions having effect to other genes in the genomic locus. For example, HiChIP has been used to delineate promoter-enhancer interactions in keratinocytes and CD8+ T-cell lines exploring psoriasis and psoriatic arthritis disease-associated SNPs and similar methods could be explored in AS. (38, 39) Additionally, targeted *RUNX3* enhancer element genomic editing strategies are also needed to elucidate the effects on RUNX3 expression and downstream cellular signaling.

In conclusion, the work presented here provides new insights into the complex transcriptional regulation of *RUNX3* and the role of AS-associated variants and the importance of interrogating their role in the appropriate cellular context.

## AUTHORS CONTRIBUTIONS

CJC, MV, JCK and BPW conceived and designed the experiments. MV and CJC performed the experiments. MV, CJC and CD analysed the data. MV, CJC, JCK and BPW drafted the manuscript, and all the authors revised the final version prior submission.

## REFERENCES

1. Bridgewood C, Sharif K, Sherlock J, Watad A, McGonagle D. Interleukin-23 pathway at the enthesis: The emerging story of enthesitis in spondyloarthropathy. Immunological reviews. 2020;294(1):27–47. Epub 2020/01/21.

2. Stolwijk C, van Tubergen A, Castillo-Ortiz JD, Boonen A. Prevalence of extra-articular manifestations in patients with ankylosing spondylitis: a systematic review and meta-analysis. Annals of the rheumatic diseases. 2015;74(1):65–73. Epub 2013/09/04.

3. Rizzo A, Ferrante A, Guggino G, Ciccia F. Gut inflammation in spondyloarthritis. Best practice & research Clinical rheumatology. 2017;31(6):863–76. Epub 2018/12/05.

4. Brewerton DA, Hart FD, Nicholls A, Caffrey M, James DC, Sturrock RD. Ankylosing spondylitis and HL-A 27. Lancet. 1973;1(7809):904–7. Epub 1973/04/28.

5. Schlosstein L, Terasaki PI, Bluestone R, Pearson CM. High association of an HL-A antigen, W27, with ankylosing spondylitis. The New England journal of medicine. 1973;288(14):704–6. Epub 1973/04/05.

6. Brown MA, Kennedy LG, MacGregor AJ, Darke C, Duncan E, Shatford JL, et al. Susceptibility to ankylosing spondylitis in twins: the role of genes, HLA, and the environment. Arthritis and rheumatism. 1997;40(10):1823–8. Epub 1997/10/23.

7. Cortes A, Pulit SL, Leo PJ, Pointon JJ, Robinson PC, Weisman MH, et al. Major histocompatibility complex associations of ankylosing spondylitis are complex and involve further epistasis with ERAP1. Nature communications. 2015;6:7146. Epub 2015/05/23.

8. Burton PR, Clayton DG, Cardon LR, Craddock N, Deloukas P, Duncanson A, et al. Association scan of 14,500 nonsynonymous SNPs in four diseases identifies autoimmunity variants. Nature genetics. 2007;39(11):1329–37. Epub 2007/10/24.

9. Reveille JD, Sims AM, Danoy P, Evans DM, Leo P, Pointon JJ, et al. Genome-wide association study of ankylosing spondylitis identifies non-MHC susceptibility loci. Nature genetics. 2010;42(2):123–7. Epub 2010/01/12.

10. Evans DM, Spencer CC, Pointon JJ, Su Z, Harvey D, Kochan G, et al. Interaction between ERAP1 and HLA-B27 in ankylosing spondylitis implicates peptide handling in the mechanism for HLA-B27 in disease susceptibility. Nature genetics. 2011;43(8):761–7. Epub 2011/07/12.

11. Cortes A, Hadler J, Pointon JP, Robinson PC, Karaderi T, Leo P, et al. Identification of multiple risk variants for ankylosing spondylitis through high-density genotyping of immune-related loci. Nature genetics. 2013;45(7):730–8. Epub 2013/06/12.

12. Ellinghaus D, Jostins L, Spain SL, Cortes A, Bethune J, Han B, et al. Analysis of five chronic inflammatory diseases identifies 27 new associations and highlights disease-specific patterns at shared loci. Nature genetics. 2016;48(5):510–8. Epub 2016/03/15.

13. Robinson PC, Leo PJ, Pointon JJ, Harris J, Cremin K, Bradbury LA, et al. The genetic associations of acute anterior uveitis and their overlap with the genetics of ankylosing spondylitis. Genes and immunity. 2016;17(1):46–51. Epub 2015/11/27.

14. Egawa T, Tillman RE, Naoe Y, Taniuchi I, Littman DR. The role of the Runx transcription factors in thymocyte differentiation and in homeostasis of naive T cells. The Journal of experimental medicine. 2007;204(8):1945–57. Epub 2007/07/25.

15. Ebihara T, Song C, Ryu SH, Plougastel-Douglas B, Yang L, Levanon D, et al. Runx3 specifies lineage commitment of innate lymphoid cells. Nature immunology. 2015;16(11): 1124–33. Epub 2015/09/29.

16. Behr FM, Chuwonpad A, Stark R, van Gisbergen K. Armed and Ready: Transcriptional Regulation of Tissue-Resident Memory CD8 T Cells. Frontiers in immunology. 2018;9:1770. Epub 2018/08/23.

17. Vecellio M, Chen L, Cohen CJ, Cortes A, Li Y, Bonham S, et al. Functional Genomic Analysis of a RUNX3 Polymorphism Associated With Ankylosing Spondylitis. Arthritis Rheumatol. 2021;73(6):980–90. Epub 2020/12/29.

18. Fishilevich S, Nudel R, Rappaport N, Hadar R, Plaschkes I, Iny Stein T, et al. GeneHancer: genome-wide integration of enhancers and target genes in GeneCards. Database : the journal of biological databases and curation. 2017;2017. Epub 2017/06/13.

19. Allen EK, Randolph AG, Bhangale T, Dogra P, Ohlson M, Oshansky CM, et al. SNP-mediated disruption of CTCF binding at the IFITM3 promoter is associated with risk of severe influenza in humans. Nature medicine. 2017;23(8):975–83. Epub 2017/07/18.

20. Schneider CA, Rasband WS, Eliceiri KW. NIH Image to ImageJ: 25 years of image analysis. Nature methods. 2012;9(7):671–5. Epub 2012/08/30.

21. Miele A, Gheldof N, Tabuchi TM, Dostie J, Dekker J. Mapping chromatin interactions by chromosome conformation capture. Current protocols in molecular biology. 2006;Chapter 21:Unit 21 11. Epub 2008/02/12.

22. Vecellio M, Cortes A, Roberts AR, Ellis J, Cohen CJ, Knight JC, et al. Evidence for a second ankylosing spondylitis-associated RUNX3 regulatory polymorphism. RMD open. 2018;4(1):e000628. Epub 2018/03/14.

23. Li Z, Haynes K, Pennisi DJ, Anderson LK, Song X, Thomas GP, et al. Epigenetic and gene expression analysis of ankylosing spondylitis-associated loci implicate immune cells and the gut in the disease pathogenesis. Genes and immunity. 2017;18(3):135–43. Epub 2017/06/18.

24. Shi C, Ray-Jones H, Ding J, Duffus K, Fu Y, Gaddi VP, et al. Chromatin Looping Links Target Genes with Genetic Risk Loci for Dermatological Traits. The Journal of investigative dermatology. 2021. Epub 2021/02/20.

25. Puig-Kroger A, Aguilera-Montilla N, Martinez-Nunez R, Dominguez-Soto A, Sanchez-Cabo F, Martin-Gayo E, et al. The novel RUNX3/p33 isoform is induced upon monocyte-derived dendritic cell maturation and downregulates IL-8 expression. Immunobiology. 2010;215(9-10):812–20. Epub 2010/07/10.

26. Estecha A, Aguilera-Montilla N, Sanchez-Mateos P, Puig-Kroger A. RUNX3 regulates intercellular adhesion molecule 3 (ICAM-3) expression during macrophage differentiation and monocyte extravasation. PloS one. 2012;7(3):e33313. Epub 2012/04/06.

27. Corbin AL, Gomez-Vazquez M, Berthold DL, Attar M, Arnold IC, Powrie FM, et al. IRF5 guides monocytes toward an inflammatory CD11c(+) macrophage phenotype and promotes intestinal inflammation. Science immunology. 2020;5(47). Epub 2020/05/24.

28. Fainaru O, Woolf E, Lotem J, Yarmus M, Brenner O, Goldenberg D, et al. Runx3 regulates mouse TGF-beta-mediated dendritic cell function and its absence results in airway inflammation. The EMBO journal. 2004;23(4):969–79. Epub 2004/02/07.

29. Selvarajan V, Osato M, Nah GSS, Yan J, Chung TH, Voon DC, et al. RUNX3 is oncogenic in natural killer/T-cell lymphoma and is transcriptionally regulated by MYC. Leukemia. 2017;31(10):2219–27. Epub 2017/01/26.

30. Lee CW, Ito K, Ito Y. Role of RUNX3 in bone morphogenetic protein signaling in colorectal cancer. Cancer research. 2010;70(10):4243–52. Epub 2010/05/06.

31. Hosoi H, Niibori-Nambu A, Nah GSS, Bahirvani AG, Mok MMH, Sanda T, et al. Superenhancers for RUNX3 are required for cell proliferation in EBV-infected B cell lines. Gene. 2021;774:145421. Epub 2021/01/15.

32. Pello OM, De Pizzol M, Mirolo M, Soucek L, Zammataro L, Amabile A, et al. Role of c-MYC in alternative activation of human macrophages and tumor-associated macrophage biology. Blood. 2012;119(2):411–21. Epub 2011/11/10.

33. Palstra RJ, Grosveld F. Transcription factor binding at enhancers: shaping a genomic regulatory landscape in flux. Frontiers in genetics. 2012;3:195. Epub 2012/10/13.

34. de Wit E, de Laat W. A decade of 3C technologies: insights into nuclear organization. Genes & development. 2012;26(1):11–24. Epub 2012/01/05.

35. Vecellio M, Roberts AR, Cohen CJ, Cortes A, Knight JC, Bowness P, et al. The genetic association of RUNX3 with ankylosing spondylitis can be explained by allele-specific effects on IRF4 recruitment that alter gene expression. Annals of the rheumatic diseases. 2016;75(8):1534–40. Epub 2015/10/11.

36. Espinola SM, Gotz M, Bellec M, Messina O, Fiche JB, Houbron C, et al. Cis-regulatory chromatin loops arise before TADs and gene activation, and are independent of cell fate during early Drosophila development. Nature genetics. 2021;53(4):477–86. Epub 2021/04/03.

37. Ing-Simmons E, Vaid R, Bing XY, Levine M, Mannervik M, Vaquerizas JM. Independence of chromatin conformation and gene regulation during Drosophila dorsoventral patterning. Nature genetics. 2021;53(4):487–99. Epub 2021/04/03.

38. Shi C, Rattray M, Barton A, Bowes J, Orozco G. Using functional genomics to advance the understanding of psoriatic arthritis. Rheumatology (Oxford). 2020;59(11):3137–46. Epub 2020/08/12.

39. Shi C, Rattray M, Orozco G. HiChIP-Peaks: a HiChIP peak calling algorithm. Bioinformatics. 2020;36(12):3625–31. Epub 2020/03/25.

